# Motor-evoked neural responses in auditory cortex are associated with improved sensitivity to self-generated sounds

**DOI:** 10.1101/2020.03.25.002485

**Authors:** Daniel Reznik, Noa Hacohen, Batel Buaron, Elana Zion-Golumbic, Roy Mukamel

## Abstract

Sensory perception is a product of complex interactions between the internal state of an organism and the physical attributes of a stimulus. One factor that modulates the internal state of the perceiving agent is voluntary movement. It has been shown across the animal kingdom that perception and sensory-evoked physiological responses are modulated depending on whether or not the stimulus is the consequence of voluntary actions. These phenomena are often attributed to motor signals sent to relevant sensory regions (efference copies), that convey information about expected upcoming sensory consequences. However, to date, there is no *direct* evidence in humans for efferent signals underlying these motor-sensory interactions. In the current study we recorded neurophysiological (using Magnetoencephalography) and behavioral responses from 16 healthy subjects performing an auditory detection task of faint tones. Tones were either generated by subjects’ voluntary button presses or occurred predictably following a visual cue. By introducing a constant temporal delay between button press/cue and tone delivery and applying source-level analysis we decoupled motor-evoked and auditory-evoked activity in auditory cortex. We show motor-related evoked-responses in auditory cortex following sound-triggering actions and preceding sound onset. Such evoked-responses were not found for button-presses that were not coupled with expected sounds. Furthermore, the amplitude of these evoked-responses corresponded with subsequent sound detection, suggesting their functional relevance to auditory processing. Our results provide first *direct* evidence for efferent signals in sensory cortex that are evoked by voluntary actions coupled with sensory consequences.

## Introduction

Voluntary actions are commonly coupled with expected sensory consequences – such as the sound of our footsteps during walking, or the tactile feedback when typing on a computer. Previous studies have demonstrated that perception of sensory stimuli is modulated by whether they are produced through voluntary actions of the perceiver or by an external source (sensory modulation; Crapse and Sommer, 2008a, b; Hughes and Waszak, 2011). For example, in the auditory domain, perceived loudness is typically attenuated for salient self-generated sounds compared to identical sounds produced by someone else (e.g., Weiss et al., 2011). Similarly, perceived tactile pressure is lower when it is applied by the perceiver vs. by an external source (Blakemore et al., 1998; Kilteni and Ehrsson, 2020; Shergill et al., 2005; Walsh et al., 2011), an effect also associated with the phenomenon that one cannot tickle oneself (Blakemore et al., 1999).

A common explanation for these phenomena at the neural level, proposed in the early 1950s, is that during voluntary execution of actions, the motor system sends an “efference copy” (Von Holst, 1954) conveying the expected sensory outcome to relevant sensory cortex. This signal results in a local “corollary discharge” (Sperry, 1950), that changes the neural state of the sensory system. Since these early studies, the role of motor cortex in modulating neural activity in sensory regions during action generation, as well as its role in perceptual modulations, has been extensively studied. Modulations have been reported both in rodents and in humans across various tasks and modalities, including auditory, visual, and tactile (Crapse and Sommer, 2008a,b; Eliades and Wang, 2003; Lee and Middlebrooks, 2011; Morillon et al., 2016; Reznik and Mukamel, 2019; Saleem et al., 2013). The results of these studies suggest that voluntary actions may play an important role in perception, by directly modulating neural activity in sensory circuits.

In the auditory domain in humans, sound-triggering actions have been shown to modulate auditory evoked responses. The magnitude of the N100/M100 component of the EEG/MEG sound-evoked response is typically attenuated when sounds are actively generated by the perceiver relative to the response evoked by identical sounds heard passively (Baess et al., 2011; Buran et al., 2014; Morillon et al., 2014; Nelson et al., 2013; Reznik et al., 2015a; Reznik et al., 2014; Schneider et al., 2014; Weiss et al., 2011; Zhou et al., 2014). However, since in these paradigms the sound is usually presented immediately following the generating action (e.g. button press), when measuring the sound-evoked signal it is difficult to disambiguate the relative contribution of action-related signals from bottom-up sensory signals evoked by the stimulus. Furthermore, in the case of EEG, motor responses and auditory responses share overlapping scalp-distributions due to the geometry of the sources relative to the scalp (Horvath, 2015), which additionally limits the ability to separate between the potential contribution of different neural sources to the observed modulations of evoked responses. Therefore, previous results are insufficient for providing direct evidence for motor-induced modulations in human auditory cortex during perception of self-generated sounds.

The current study was aimed at providing this missing link. We examined how sound-eliciting button presses affect (1) the neural state in auditory cortex *preceding* sound onset, (2) evoked responses *following* the sound, and (3) perception of low amplitude sounds. To this end, subjects performed a sound detection task of faint auditory stimuli, that were physically identical but either self-generated (Active condition via button presses) or generated by the computer (Passive condition). Critically, in the active condition, we introduced a constant temporal delay between the action and its auditory outcome in order to dissociate bottom-up sound-locked sensory responses from action-locked neural modulations in auditory cortex. Neural activity was measured using Magnetoencephalography (MEG) and analyzed in source space, further allowing us to distinguish between neural responses in motor and auditory cortices. Our results provide direct evidence for action-evoked activity in auditory cortex, which is also associated with perceptual modulations of self-generated sounds.

## Results

### Behavioral data

After assessing individual auditory thresholds, each subject was engaged in a sound-detection task in which faint sounds were presented on 50% of the trials (see STAR methods). Mean sound detection rate across subjects was 0.58±0.04 on trials in which sounds were presented (hits). In agreement with our previous reports (Reznik et al., 2015a; Reznik et al., 2014), participants showed increased sensitivity to sounds in the Active compared to Passive condition (mean±SEM *d’* across subjects: Active *d’*=2.07±0.19, Passive *d’*=1.82±0.19; paired one-tailed t-test, t_(15)_=2.28, p=0.03; see Figure 1b for group and individual subjects data). Importantly, participants showed no difference in bias for responding “yes” vs. “no” sound detection across conditions (mean±SEM *c* across subjects: Active *c*=0.74±0.05, Passive *c*=0.75±0.06; paired two-tailed t-test, t_(15)_=0.13, p=0.89; Figure 1b). In the active condition, we did not impose a temporal limit between the appearance of the visual cue and subjects’ button presses. Subjects responded on average 936±35ms following the visual cue with a non-significant tendency for faster responses in detected vs. non-detected (hit vs. miss) trials (mean±SEM response time across subjects: Active-hit: 0.90±0.04ms, Active-miss: 1.00±0.06ms; two-tailed paired t-test, t_(15)_=1.76, p=0.09).

**Figure 1.**
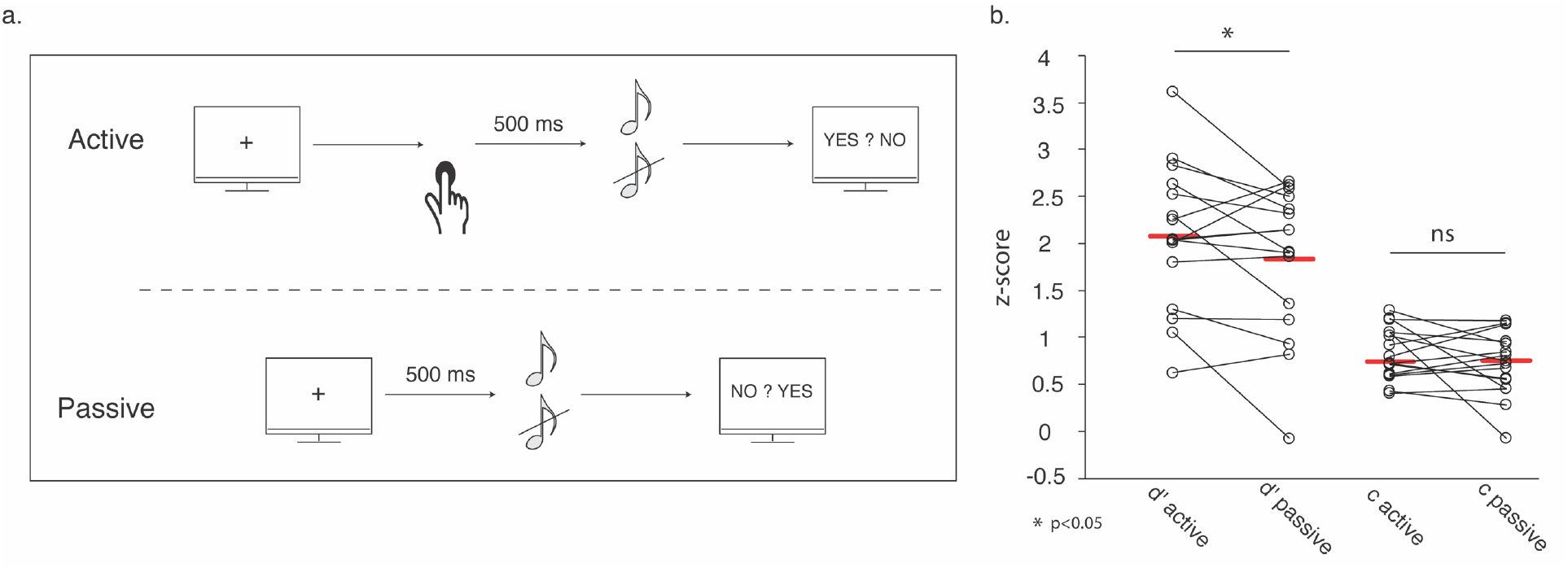
Experimental design and performance in the auditory detection task. (**a**) In the active condition, subjects where visually cued to perform a button press with their right index finger. Subjects pressed the button at their own time and 500ms following the button press a pure tone was delivered in 50% of the trials. In the passive condition, a pure tone was delivered in 50% of the trials 500ms following a visual cue. At the end of each trial, subjects answered whether they heard the tone by pressing a button with their left hand. In both conditions, tones were delivered at individual subjects’ auditory detection levels. (**b**) Group-level (n=16) comparisons of performance in the auditory detection task. Group means are shown with red lines and individual subjects are shown with dots. While subjects’ auditory sensitivity (measured with *d’*) was greater in the active compared with passive conditions, subjects’ criterion did not differ between the two conditions.

In addition to the active and passive conditions, subjects also performed a Motor-only task in which they pressed buttons with no auditory consequences (see Methods). In this task, response times to the visual “+” cue were faster than in the Active trials, averaging 582±59ms (n=14; due to technical reasons, response times in this condition were not measured in 2 subjects; t_(13)_=5.81, p<0.001).

### Pre-sound, neural activity in auditory cortex

First, we compared neural activity in the Active vs. Passive conditions during the 500-ms window following the sound-triggering button press (Active) or visual cue indicating an upcoming sound (Passive) and until the onset of the sound itself. This analysis focused on MEG activity in bilateral auditory cortices (left/right ROIs). We performed a 2×2 repeated measures ANOVA with the factors of Condition (Active/Passive) and Hemisphere (right/left). This 2×2 analysis revealed a main effect of Condition (F_(1,15)_=11.89; p=0.004), main effect of Hemisphere (F_(1,15)_=14.08; p=0.002) and a significant interaction effect (F_(1,15)_=6.11; p=0.026). Post-hoc pair-wise comparisons indicated that while MEG signal was greater in the Active compared with the Passive conditions in the left auditory cortex (mean±SEM dSPM across subjects - Active: 2.31±0.14, Passive: 1.68±0.09, t_(15)_=4.21, p_holm_<0.01), no difference was observed in the right hemisphere (Active: 1.64±0.13, Passive: 1.52±0.10, t_(15)_=0.84, p_holm_=0.85; Figure 2).

**Figure 2.**
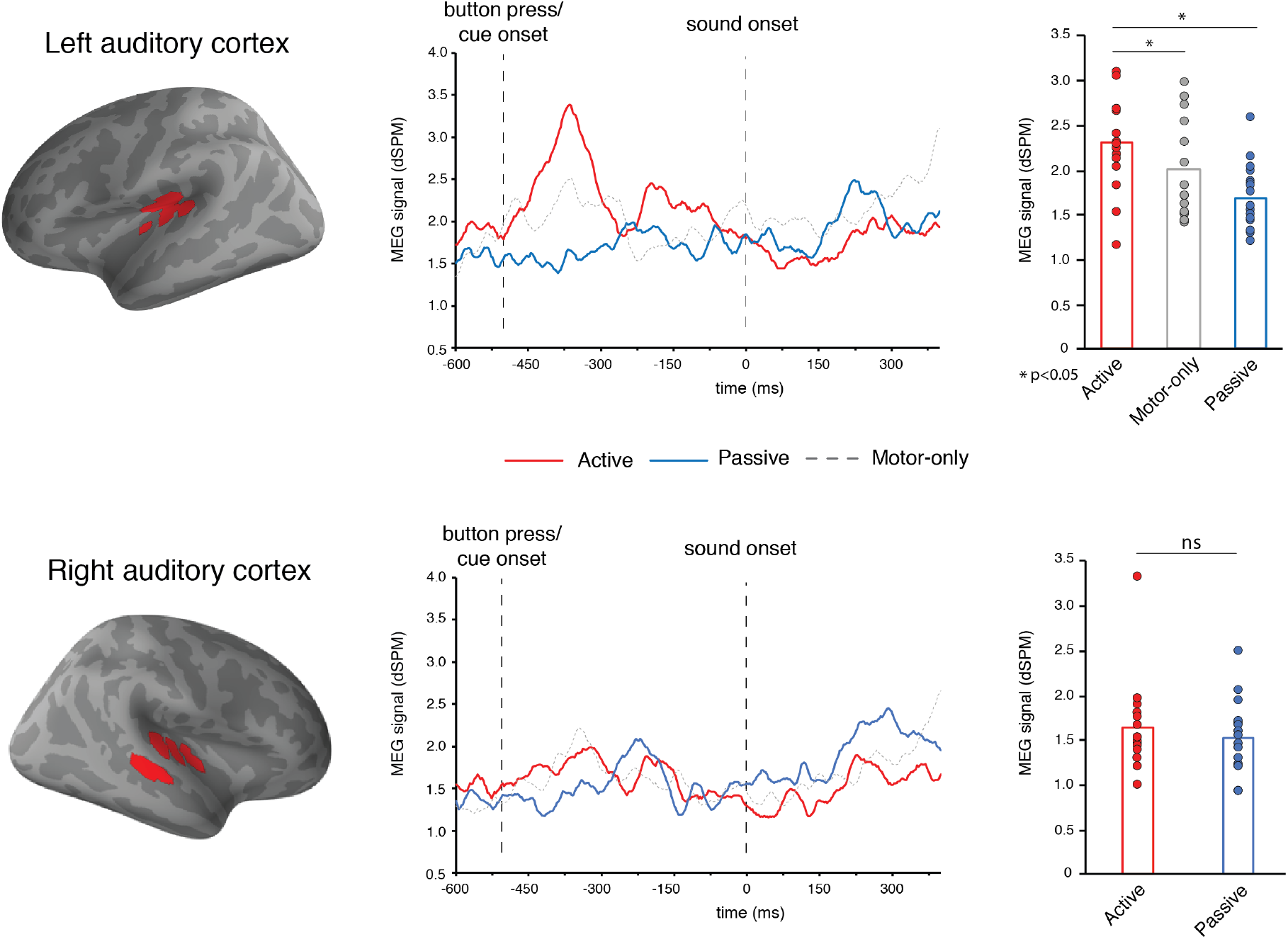
MEG time-course in auditory cortex during active, passive and motor-only conditions. **Left panel**: left (top) and right (bottom) auditory ROIs defined on the group level (n=16) and projected to the cortical surface template. **Middle panel**: group average (n=16) time-courses from the right and left auditory ROIs in the active, passive and motor-only conditions. **Right panel**: group-level (n=16) comparisons of the MEG signal in the 500ms pre-sound time-window. Group means are shown with bars and individual subjects are shown with dots.

In principle, the enhanced activity we find in left auditory cortex in the active condition during the pre-sound period could be related to button presses irrespective of their associated auditory outcome. To test this possibility, we compared this response to activity during the same time-window in the Motor-only condition (i.e. button presses with no auditory consequences). Indeed, activity in left auditory cortex during the Active condition was significantly higher than in the Motor-only condition in which no sound was expected or delivered (mean±SEM dSPM across subjects – Motor-only: 2.01±0.14, two-tailed paired t-test, t_(15)_=2.36, p=0.03; Figure 2). Responses in the Motor-only condition during this time-window had a tendency to be higher than the Passive condition, but this did not reach significance (t_(15)_=1.90, p=0.08). Taken together, these results suggest that the enhanced activity in left auditory cortex during Active compared with Passive condition is not the mere result of right hand finger presses but rather the association of these button presses with an expected auditory consequence.

Next, we tested whether activity in left auditory cortex during the pre-sound period was associated with subsequent sound perception, in both Active and Passive conditions. Specifically, we examined whether neural responses differed between trials in which subjects detected vs. failed to detect the threshold-level sounds (‘hit’ vs. ‘miss’ trials) in each condition. To this end, we performed a 2×2 repeated measures ANOVA with the factors of Condition (Active/Passive) and Detection (hit/miss). We found a main effect of condition (F_(1,15)_=19.87; p<0.001), with stronger responses in the Active vs. Passive condition (as before), as well as a main effect of detection (F_(1,15)_=6.70; p=0.02; Figure 3) indicating that the MEG signal in the period preceding the sound was greater in ‘hit’ vs. ‘miss’ trials. The interaction between the factors was not significant (F_(1,15)_=0.02; p=0.91; Figure 3b). The same analysis performed in the right hemisphere yielded no significant results (Condition effect - F_(1,15)_=0.683; p=0.42; Detection effect - F_(1,15)_=0.37; p=0.55; Interaction effect - F_(1,15)_=3.13; p=0.09; Figure 3b).

**Figure 3.**
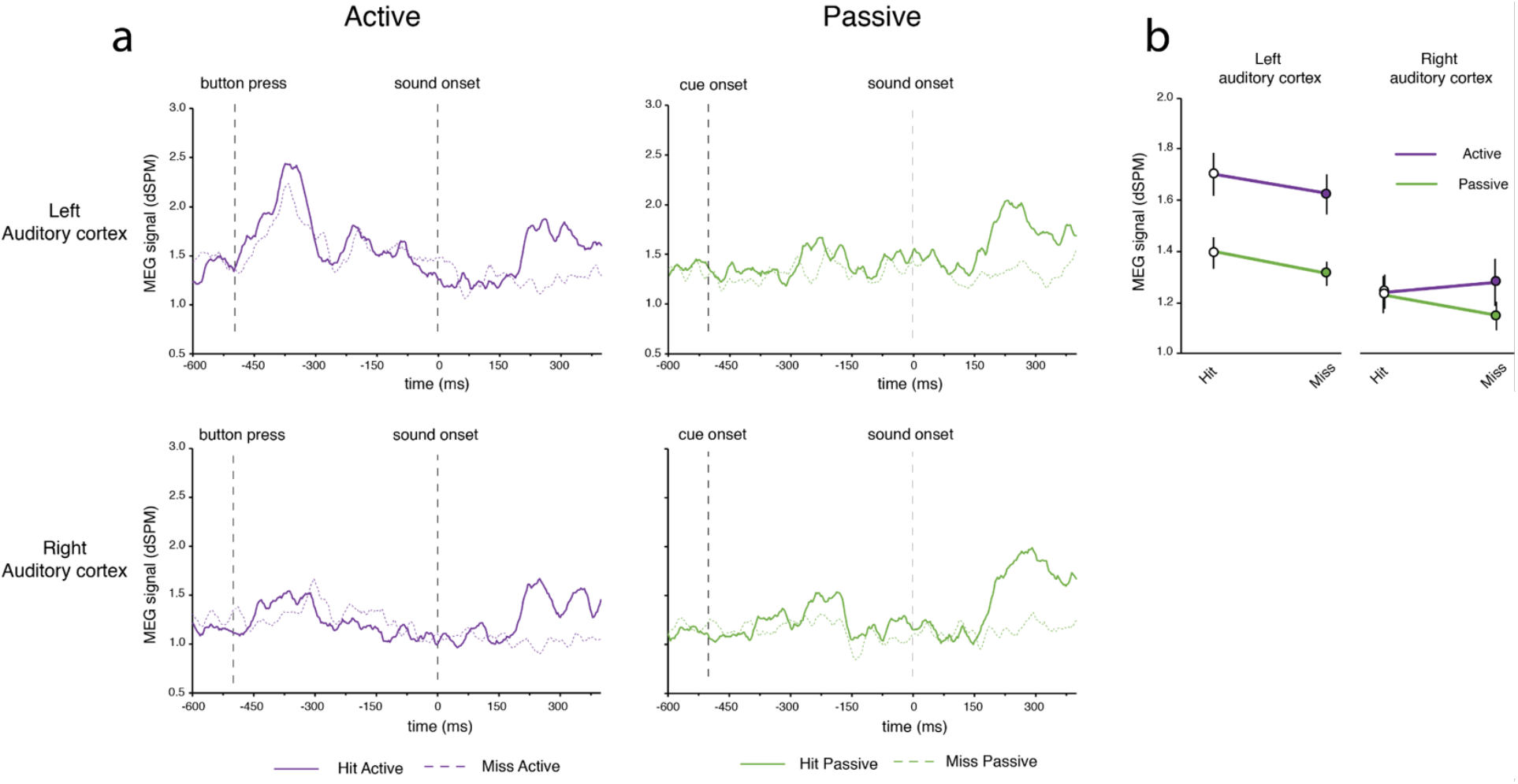
MEG time-course in auditory cortex during hit and miss trials in active and passive conditions. (**a**) Group average (n=16) time-courses from the left (top) and right (bottom) auditory ROIs in hit and miss active/passive trials (**b**) Group-level (n=16) analyses of variance of the MEG signal in the 500ms pre-sound time-window. Group means are shown with circles and standard errors with vertical bars. In the left auditory cortex, MEG activity was greater prior to hit compared with miss trials.

Visual inspection of the time-course in the pre-sound period showed a peak centered ~140 ms following the button press in the Active condition (~360ms prior to sound onset). A post-hoc analysis, following up on the observed difference between activity in left auditory cortex preceding ‘hits’ vs. ‘misses’, focused on paired comparison in a shorter 100ms time window around this peak. This analysis confirmed the greater response prior to ‘hit’ compared with ‘miss’ trials in the Active condition (mean±SEM dSPM across subjects – Active hit: 2.23±0.19, Active miss: 1.96±0.16, two-tailed paired t-test, t_(15)_=2.36, p=0.03), but no significant effect of subsequent performance was found in the Passive condition in the corresponding time window (Passive hit: 1.33±0.07, Passive miss: 1.26±0.05, t_(15)_=0.9, p=0.37).

### Sound-evoked neural activity in auditory cortex

Analysis of the evoked-responses in auditory cortex to the threshold-level sounds focused on a 100ms time widow centered around the peak of the group evoked response collapsed across Active and Passive conditions (230ms following sound onset; Figure 2). This window was then used to extract the corresponding data of individual subjects. We performed a 2×2 repeated measures ANOVA with the factors Condition (Active/Passive) and Hemisphere (right/left). This 2×2 analysis yielded a main effect of Condition (F_(1,15)_=5.47; p=0.034), no main effect of Hemisphere (F_(1,15)_=0.48; p=0.51) and no interaction effect (F_(1,15)_=0.084; p=0.78). Planned paired-comparisons of these effects separately in each hemisphere, confirmed a weaker evoked responses in both hemispheres in the Active compared with Passive conditions (mean±SEM dSPM across subjects, Left auditory cortex - Active: 1.81±0.21, Passive: 2.29±0.27, one-tailed paired t-test, t_(15)_=2.16, p=0.04; Right auditory cortex - Active: 1.71±0.13, Passive: 2.21±0.22, t_(15)_=3.11, p =0.007; Figure 4a).

**Figure 4.**
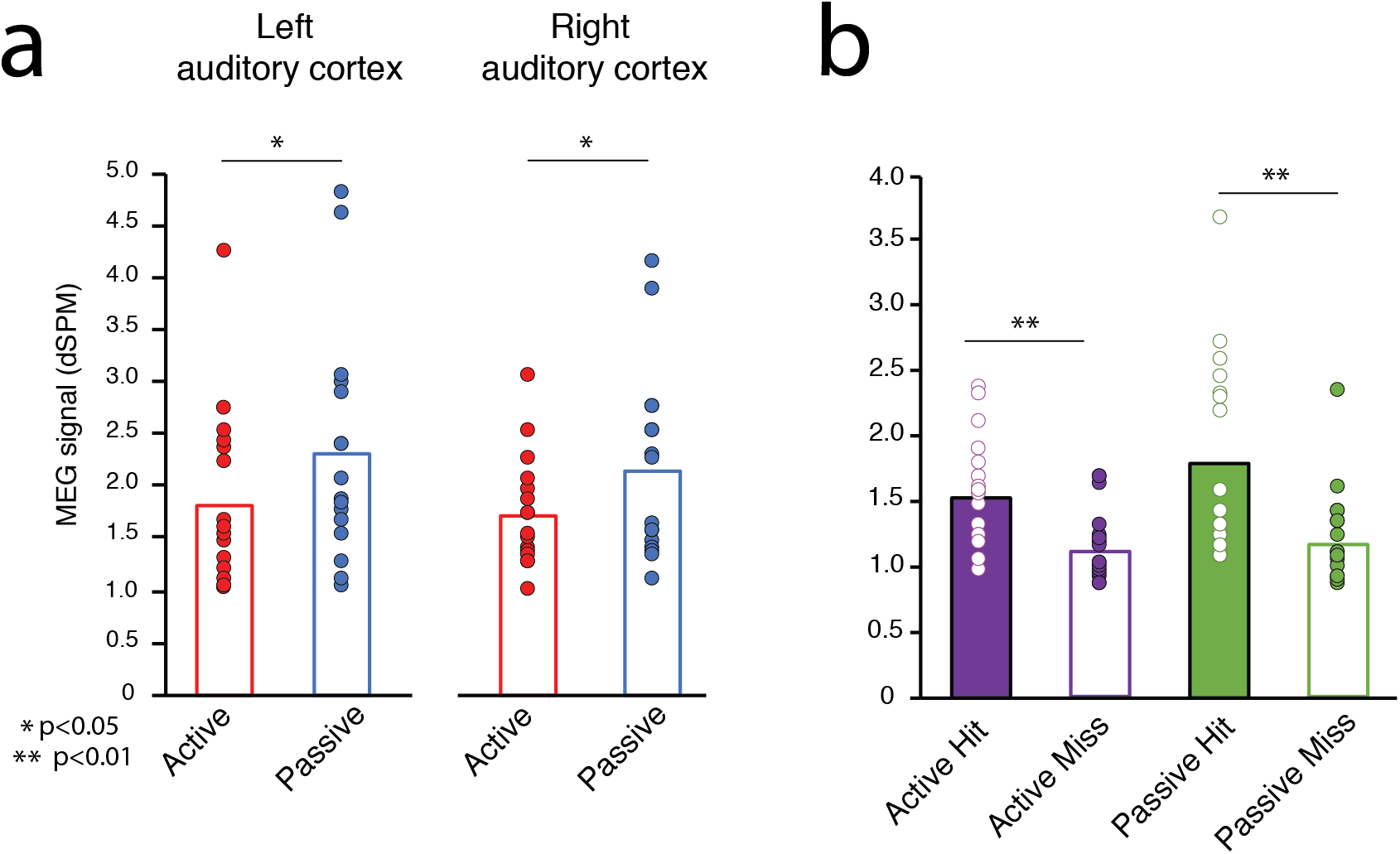
Sound-evoked activity in auditory cortex. (**a**) Group-level (n=16) comparisons of sound-evoked activity in right and left auditory cortex. Sound-evoked activity was reduced in active compared with passive conditions in right and left auditory cortex (see also post-sound time course in Figure 3). (**b**) Group-level (n=16) comparisons of sound-evoked activity in hit and miss trials in auditory cortex. In both conditions (active/passive), sound-evoked activity was greater when subjects successfully detected the auditory tones (hit trials) compared with when subjects failed to do so (miss trials; collapsed across right and left auditory cortices). Group means are shown with bars and individual subjects are shown with dots.

Similar to the analysis we performed in the pre-sound epoch, we examined whether the magnitude of the evoked-response to the sounds was associated with subjective perception. To this end we compared the auditory evoked responses during ‘hit’ and ‘miss’ trials in Active and Passive conditions in right and left auditory cortices. We performed a 2×2×2 repeated measures ANOVA with the factors Condition (Active/Passive), Hemisphere (right/left) and Detection (hit/miss). This analysis yielded only a main effect of Detection (F_(1,15)_=17.05; p=0.001). Planned paired comparisons of the Detection effect collapsed across hemispheres showed that activity in auditory cortex during “hit” trials was greater than the activity in “miss” trials (mean±SEM dSPM across subjects – Active Hit: 1.54±0.10; Active Miss: 1.13±0.07; two-tailed paired t-test, t_(15)_=4.16, p<0.001; Passive Hit: 1.79±0.18; Passive Miss: 1.18±0.09; two-tailed paired t-test, t_(15)_=4.26, p<0.001; Figure 4b). No other main effects were significant (all p>0.074).

### Action-locked neural activity in motor cortex

Although our primary research questions pertained to the modulatory effects of sound-triggering actions on neural activity in auditory cortex, the whole-brain coverage of the MEG allowed us to also explore whether activity in motor cortex is modulated by expected auditory consequences. To this end, we compared activity in motor cortex evoked by mere (silent) button presses in the Motor-only condition vs. button presses in the Active condition that generated sounds on 50% of the trials. Focusing on a 500ms time window following the button press, we found that activity in somatomotor cortex was enhanced in the Active condition relative to the Motor-only condition (mean±SEM dSPM across subjects – Active: 3.36±0.41, Motor-only: 2.79±0.40; two-tailed paired t-test, t_(15)_=2.33, p=0.03; Figure 5). Visual inspection of the time-course shows that in the Active condition, activity in motor cortex is sustained throughout the entire 500ms window, suggesting that motor cortex remains engaged in anticipation of the upcoming coupled sound. No differences in motor cortex activity were found between Active Hit vs Active Miss trials (mean±SEM dSPM across subjects – Active Hit: 2.47±0.41; Active Miss: 2.58±0.53; two-tailed paired t-test, t_(15)_=0.77, p=0.45).

**Figure 5.**
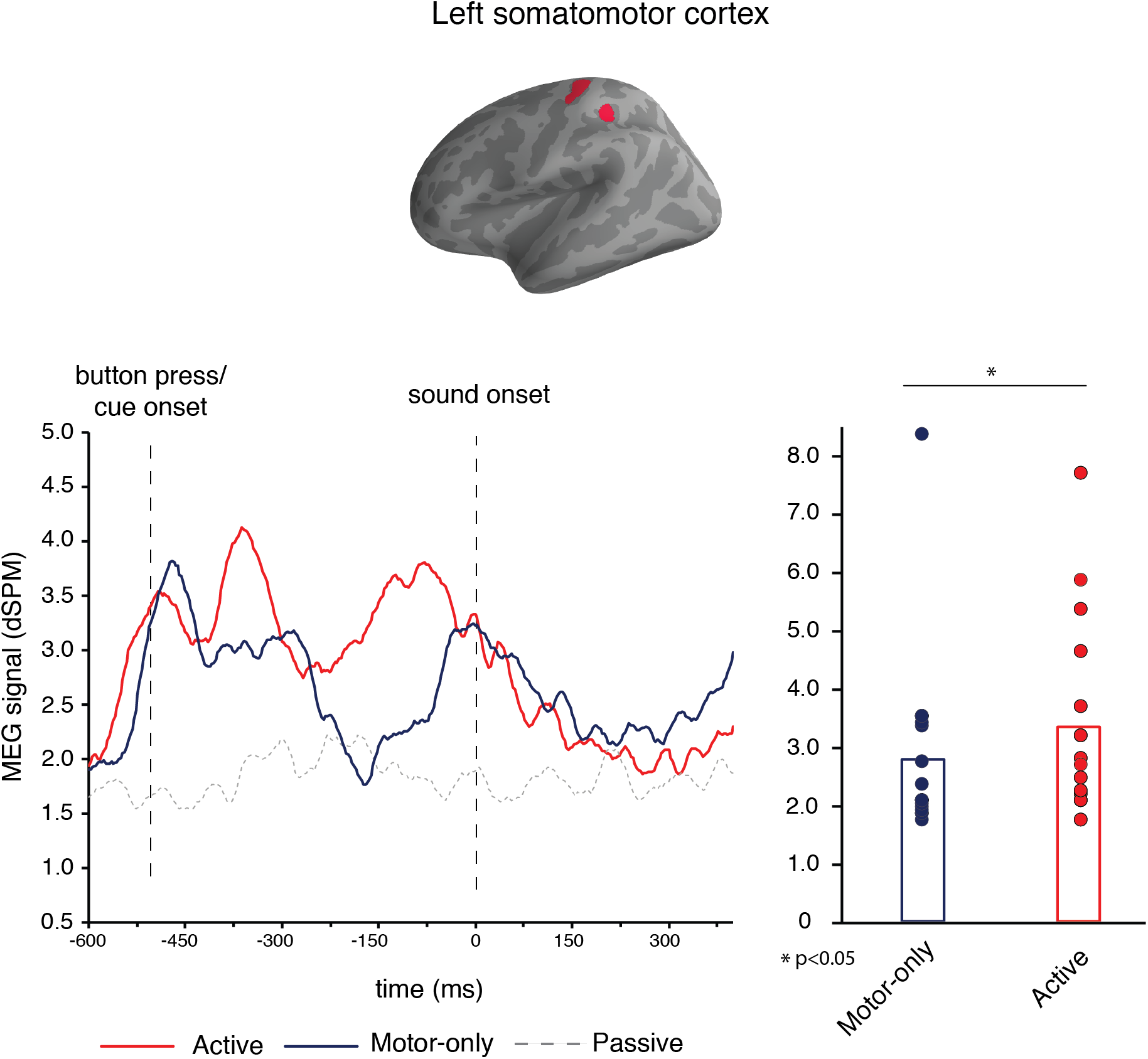
Somatomotor activity following button press. **Top panel**: left somatomotor ROI defined at the group level (n=16) and projected to the cortical surface template. **Bottom left panel**: group average (n=16) time-courses from the somatomotor ROIs in the active, passive and motor-only conditions. **Bottom right panel**: group-level (n=16) comparisons of the MEG signal in the motor-only and active conditions during the 500ms pre-sound time-window. During the 500ms time-window following button press, activity in the left somatomotor cortex was greater when button presses were associated with auditory consequences (active condition) compared with when button presses were not associated with auditory consequences (motor-only conditions). Group means are shown with bars and individual subjects are shown with dots.

## Discussion

In the current study, we examined how actions that are coupled with the generation of faint auditory sounds influence auditory perception and neural activity in auditory cortex. Using MEG, we provide evidence for motor evoked responses in auditory cortex contralateral to the active hand, an effect also associated with increased perceptual salience for weak self-generated sounds.

### Motor-evoked neural activity in auditory cortex

Although modulations of sensory-evoked responses to self-generated sounds in auditory cortex are assumed to be a consequence of input from motor regions (efference copy), there is no direct evidence for this in humans. In previous studies, sensory consequences immediately followed the action and analysis focused on the evoked response – precluding the ability to assess the independent effect of the motor actions themselves on neural activity in auditory cortex (Horvath, 2015; Saupe et al., 2013; Timm et al., 2013). A unique feature of our experimental design was the introduction of a temporal delay between the motor act and its auditory consequence. This feature, together with the high spatial and temporal resolution of MEG, allowed us to detect neural signals within auditory cortex that follow the motor act and are not yet confounded by bottom-up sound-evoked activity. In left auditory cortex we found a significant increase in MEG signal locked to button press. This increase was not seen when subjects passively heard identical sounds, and was significantly weaker when subjects only pressed buttons with no expectation for auditory consequences. These results are consistent with the notion of an efference-copy modulating neural states in sensory cortex during perception of expected self-generated sensory consequences.

Interestingly, the action-locked increase in MEG signal we report in left auditory cortex was not found in right auditory cortex (ipsilateral to the button-pressing hand). This is compatible with the fact that subjects used their right index finger to press the buttons. In a previous fMRI study we found stronger modulations in auditory cortex for sounds produced with the contra-lateral hand (or ipsi-lateral to the active motor cortex; (Reznik et al., 2014; Reznik et al., 2015b). Recently we examined whether this limb-specificity is also found in other modalities and find that fMRI signals in visual cortex are differentially modulated according to the hand (right/left) that was used to elicit the visual stimulus (Buaron et al., 2020). Taken together, current MEG data further support the notion that modulations in sensory regions have a component of limb-specificity.

In principle, the evoked response we find in left auditory cortex following the button press could be due to tactile feedback rather than motor ‘efference’ copy signaling the upcoming auditory outcome of the action. Previous studies in monkeys, and humans report tactile-evoked responses in auditory cortex (Foxe et al., 2000; Foxe et al., 2002; Schroeder et al., 2001). However, we note that similar tactile feedback during button presses without auditory consequences (our Motor-Only condition), elicited motor-evoked signals in auditory cortex that were significantly reduced. Therefore, although we cannot rule out some tactile component in the motor-evoked signals in Active trials, such a component cannot fully explain the result (see also Reznik et al., 2015a).

In the human literature, the term sensory attenuation has been used to describe both the behavioral phenomenon of reduced perception (e.g. level of ticklishness, or sound loudness), and the physiological phenomenon of decreased evoked responses measured by EEG or MEG (N100 or M100, respectively). However, to date, there is no direct causal evidence linking the two phenomena. Moreover, a recent study by Palmer and colleagues even suggests different underlying mechanisms (Palmer et al., 2016). Our results demonstrate that motor-evoked signals in auditory cortex are stronger for sounds that were detected (hits) compared with sounds that were not (misses). Although still correlative evidence at this stage, these results suggest that action-induced neural modulations in auditory cortex may play a role in the behavioral manifestation of sensory modulations. The causal link between such motor-evoked physiological markers in auditory cortex and behavioral reports requires further studies.

### Sound-evoked neural activity in auditory cortex

With respect to sound-evoked neural responses in auditory cortex, we find correspondence between perception (hit/miss) and magnitude of the evoked responses. Since we used faint sounds with fixed amplitude at the subject’s hearing threshold, their detection varied across trials. We find that sound-evoked responses in trials in which the sound was perceived (hit trials) were stronger than the sound-evoked responses of trials in which sounds were not detected (miss trials). These results are in agreement with previous studies reporting physiological correlations with behavior (Hillyard et al., 1971; Jones et al., 2007). With respect to time-to-peak of the evoked response, we note that in our current results it was late (~230ms at the group level) relative to typical auditory evoked responses (~100ms). This might be due to the nature of our faint auditory stimuli. Indeed, there is evidence for a dependency of peak latency on sound intensity, with lower sound intensities associated with longer N100 latency (Neukirch et al., 2002).

### Behavior

Our behavioral results point to increased perceptual sensitivity in the active vs. passive condition. Previous studies examining sensory modulations in the auditory domain have used comparison tasks in which subjects report relative subjective loudness of salient sounds generated by self or other (e.g. Weiss et al., 2011). In a previous study, we manipulated the intensity of the expected sound and reported sensory attenuation for self-generated loud sounds (in agreement with earlier literature), and sensory enhancement when expected sounds were weak (Reznik et al., 2015a). In the current study we opted to use low amplitude sounds that allowed us to use a detection task rather than a comparison task of two supra-threshold sounds as in earlier studies. In alignment with previous findings, we report that sound detection as assessed by d’ measures, was better when faint sounds were the consequence of button presses rather than externally cued (Reznik et al., 2015a; Reznik et al., 2014).

Our motivation for introducing a temporal delay between the action and its auditory consequences was to disambiguate neural signals locked to the action from signals evoked by the sounds. However, this design also provides important information with respect to the temporal window in which action-induced modulations affect perception. In the tactile domain, behavioral results show that a delay of ~300ms abolishes perceptual attenuation (Blakemore 1999; Bays 2005). In the current study, we find that perceptual modulations for self-generated sounds are maintained even when a temporal delay of 500ms was introduced (at least in the current case where subjects were explicitly made aware of this delay). Nevertheless, characterizing the temporal window in which actions exert modulations on perceived consequences and its relation to sense of agentic control are important issues for further research (Haggard, 2017).

### Motor-evoked neural activity in motor cortex

Finally, the spatial resolution of the MEG signal allowed us to also examine neural activity in motor cortex. Although not the focus of our current study, we find that in the active condition, neural activity in left motor cortex is sustained and remains elevated throughout the intervening period between button press and sound presentation. When button presses were not associated with an auditory consequence (Motor-only condition), the MEG signal gradually returned to baseline, to rise again around 500ms later. Throughout the experiment, subjects associated button presses with the presentation of faint sounds. Since our motor-only condition was performed last during the session, we believe this late signal rise could be explained by implicit expectation of an impending sound despite the fact that the subject’s knew that no sound will be delivered. Since even in the Active condition, button-presses elicited sounds only on 50% of trials, the interesting phenomenon of sustained activity in motor cortex needs to be addressed separately by comparing actions that are either completely coupled or completely decoupled from subsequent sounds.

In summary, we show motor evoked responses in auditory cortex contralateral to the active hand, and that actions with weak auditory consequences are associated with increased perceptual salience relative to otherwise identical sounds. Our results constitute an important step in elucidating the neural mechanisms of sensory modulations and their link to perception.

## Acknowledgements

Authors declare no conflict of interest. We thank Yuval Harpaz, Adi Korisky, Maor Wolf, Uri Berger and Ohad Felsenstein for their help with MEG protocols, Shahar Moskovich for help with data collection and behavioral data analysis. This research was supported by The Israel Science Foundation (grant No. 2392/19) to R.M.; Sagol School of Neuroscience and the Israeli Presidential Honorary Scholarship for Neuroscience Research to D.R.

## Author Contributions

D.R., E.Z.G. and R.M. conceived the study and the experimental design; D.R. and N.H. collected the data; D.R., N.H. and B.B. analyzed the data; D.R., N.H., B.B., E.Z.G. and R.M. wrote the manuscript.

## STAR Methods

### Participants

Eighteen participants naïve to the purpose of the experiment were recruited to this study. Two participants did not complete the experimental procedure (one participant withdrew in the middle and data acquisition from the other participant was interrupted by a fire-alarm in the MEG facility), yielding a total of 16 participants (11 females; mean age – 22.9, range – 19 to 27 years) for subsequent analyses. All participants reported right hand dominance and normal vision and hearing. The study was approved by the Ethics committee of Bar-Ilan University and the Ethics committee of Tel-Aviv University. All participants provided written informed consent to participate in the study and were compensated for their time.

### Procedure and stimuli

Prior to the experimental task, we assessed individual participants’ auditory detection thresholds for binaural delivery of pure tones (pitch: 1 kHz, duration: 300 ms including linear rise/decay time of 25 ms) using a ‘1 step up, 2 steps down’ staircase procedure (Gelfand, 2010; 1 dB step size). On each trial, participants pressed a button with their right index finger that triggered the presentation of a tone. Participants indicated whether or not they heard it by pressing one of two buttons using their left hand. If a sound was detected, Sound Pressure Level (SPL) on the next trial was lowered by 2dB and if not, SPL on the next trial was increased by 1dB. The lowest intensity at which a subject reported detection twice (once going up and once going down) was determined as the auditory detection threshold (Gelfand, 2010). The sound-intensity corresponding to each individual’s auditory detection threshold was used throughout the main experiment, described next.

The main experiment was an auditory detection task comprised of *Active* and *Passive* experimental conditions (Figure 1). In the *Active* condition, participants saw a visual cue (“+” on a black screen) which served as a signal to press a button with their right index finger (no speed limit was imposed). In 50% of the trials, the near-threshold sound was presented 500 ms after the button press, whereas in the remaining trials no sound was delivered. Similarly, in the *Passive* condition, the same visual cue was presented on the screen but instead of cueing the subject to press the button, it indicated that a sound might be presented 500ms later (sounds occurred in 50% of the trials, similar to the active condition). In both conditions participants were asked to report whether or not they heard a sound via button press using their left hand (index/middle finger; the mapping of “yes/no” detection responses to specific fingers was randomized in each trial to avoid motor preparation). The order of sound/no sound trials was randomly mixed within each block. This design allowed us to measure the hit and false alarm rates and to calculate the sensitivity (*d’*) and criterion (*c*) measures (Green and Swets, 1974), for both active and passive conditions. During the inter-trial-interval (randomly varied between 1.5 and 2.5 s), a blank black screen was shown. The conditions were presented in 30 randomized blocks (total of 15 active and 15 passive blocks), each one consisting of 10 trials (Reznik et al., 2014). Before each block, participants were visually informed about the upcoming condition type by the words “ACTIVE” or “PASSIVE” appearing on the screen. Each block started by the subject pressing a button with their left hand.

To allow functional source localization (see below), following the main auditory detection task, participants underwent an auditory localizer task in which they passively listened to 75 salient pure tones (300ms, 1kHz) delivered binaurally with random inter-stimulus-interval ranging from 1.5 to 2.5 seconds.

Finally, to measure activity in auditory cortex associated with button presses that are not associated with auditory outcome (i.e. without sound delivery), subjects performed a silent motor task (‘Motor-only’) in which they were prompted by a visual cue (“+” on a black screen) to spontaneously perform a single button press with their right index finger. This task is similar to the Active condition blocks however the subjects were explicitly informed that their actions are no longer associated with sound delivery.

### MEG recording

During the experimental sessions, participants laid in a supine position in a magnetically shielded room while brain activity was recorded using a whole head 248-channel magnetometer array (4-D Neuroimaging, Magnes 3600 WH) located at Gonda Brain Research Center in Bar-Ilan University. Neuromagnetic activity was measured with a sampling rate of 1017.23 Hz and filtered online using 0.1-400 Hz band-pass filter. Five magnetic coils were attached to participants’ scalp to track the position of their head relative to the magnetometer array. The locations of the coils were determined with respect to 3 anatomical landmarks – nasion and left and right mastoid points. The head shape of each participant was digitized using a digital stylus (Polhemus). Sounds were delivered from a computer via Roland Octa Capture sound card through MEG-compatible in-ear earphones (Etymotic Research ER30). Subject button presses were registered using an MEG-compatible responses box (Current Designs Finger Tapper).

### MEG Pre-processing

MEG data were preprocessed using MNE toolbox for python (Gramfort et al., 2013). External noises (power-line, mechanical vibrations of the building) were removed offline using a predesigned algorithm (Tal and Abeles, 2013). A single noisy channel was excluded from analysis based on visual inspection, and signal associated with heart beats was removed using signal space projection algorithm (Uusitalo and Ilmoniemi, 1997). Data from the main experiment was segmented into 2 seconds-long epochs – from 1 second before sound onset until 1 second after sound onset. In the motor and auditory localizer tasks, data was similarly segmented into 2 seconds epochs – spanning 1 second before and after the button press or sound onset, respectively. Each trial was baseline corrected for its global mean and the average response was calculated for each condition and each subject. Since the number of trials across conditions was not identical (due to differences in the number of detected/not-detected trials, and discarded trials following signal pre-processing), we equated the number of trials within each subject across conditions by randomly selecting trials from the conditions with higher number of trials.

### MEG source localization

For each participant, we fitted the digitized head shape to a template MRI model, allowing a common source space for grand averaging across participants. MNE dynamic statistical parametric mapping (dSPM) was used to project MEG data from the sensor space to the cortical surface template. This resulted in 10,242 source reconstructed MEG signals (i.e., vertices) per hemisphere. Empirical estimate of the noise covariance was calculated from the pre-stimulus period in the auditory localizer task (200 ms window prior to sound onset), using an automated model selection procedure (Engemann and Gramfort, 2015). We used a boundary-element model (BEM) method to model activity at each vertex and calculate a forward solution. An inverse solution was estimated using this forward model and the noise estimate (SNR = 1).

### MEG data analysis

We defined three Regions of interest (ROI) – right and left auditory cortex, and left somatomotor cortex. The somatomotor ROI was identified based on activity between 100-200ms after the right hand button press in the Active and Motor-only conditions (see cortical maps in Figures 2 and 5). We used data from both conditions in order not to bias our selection of vertices when examining differences between these two conditions in left somatomotor cortex (see below). Auditory ROIs were identified based on activity in a 300ms long window following tone presentation in the auditory localizer task. ROIs (auditory and motor) were defined as all vertices in a radius of 5mm around the top 0.5 percent of active vertices in the above mentioned time-windows.

## References

Baess, P., Horvath, J., Jacobsen, T., and Schroger, E. (2011). Selective suppression of self-initiated sounds in an auditory stream: An ERP study. Psychophysiology 48, 1276–1283, 10.1111/j.1469-8986.2011.01196.x.

Bays, P. M., Wolpert, D. M., & Flanagan, J. R. (2005). Perception of the consequences of self-action is temporally tuned and event driven. Current Biology, 15(12), 1125–1128, 10.1016/j.cub.2005.05.023

Blakemore, S.J., Frith, C.D., and Wolpert, D.M. (1999). Spatio-temporal prediction modulates the perception of self-produced stimuli. J Cogn Neurosci 11, 551–559, 10.1162/089892999563607.

Blakemore, S.J., Wolpert, D.M., and Frith, C.D. (1998). Central cancellation of self-produced tickle sensation. Nat Neurosci 1, 635–640, 10.1038/2870.

Buaron, B., Reznik, D., Gilron, R., and Mukamel, R. (2020). Voluntary actions modulate perception and neural representation of action-consequences in a hand-dependent manner. bioRxiv, 10.1101/2020.01.12.903054.

Buran, B.N., von Trapp, G., and Sanes, D.H. (2014). Behaviorally gated reduction of spontaneous discharge can improve detection thresholds in auditory cortex. J Neurosci 34, 4076–4081, 10.1523/JNEUROSCI.4825-13.2014.

Crapse, T.B., and Sommer, M.A. (2008a). Corollary discharge across the animal kingdom. Nat Rev Neurosci 9, 587–600, 10.1038/nrn2457.

Crapse, T.B., and Sommer, M.A. (2008b). Corollary discharge circuits in the primate brain. Curr Opin Neurobiol 18, 552–557, 10.1016/j.conb.2008.09.017.

Eliades, S.J., and Wang, X. (2003). Sensory-motor interaction in the primate auditory cortex during self-initiated vocalizations. J Neurophysiol 89, 2194–2207, 10.1152/jn.00627.2002.

Engemann, D.A., and Gramfort, A. (2015). Automated model selection in covariance estimation and spatial whitening of MEG and EEG signals. NeuroImage 108, 328–342, 10.1016/j.neuroimage.2014.12.040.

Foxe, J.J., Morocz, I.A., Murray, M.M., Higgins, B.A., Javitt, D.C., and Schroeder, C.E. (2000). Multisensory auditory-somatosensory interactions in early cortical processing revealed by high-density electrical mapping. Cognitive Brain Research 10, 77–83, 10.1016/S0926-6410(00)00024-0.

Foxe, J.J., Wylie, G.R., Martinez, A., Schroeder, C.E., Javitt, D.C., Guilfoyle, D., Ritter, W., and Murray, M.M. (2002). Auditory-somatosensory multisensory processing in auditory association cortex: an fMRI study. Journal of neurophysiology 88, 540–543, 10.1152/jn.2002.88.1.540.

Gelfand, S. (2010). Essentials of Audiology.

Gramfort, A., Luessi, M., Larson, E., Engemann, D.A., Strohmeier, D., Brodbeck, C., Goj, R., Jas, M., Brooks, T., and Parkkonen, L. (2013). MEG and EEG data analysis with MNE-Python. Frontiers in neuroscience 7, 267, 10.3389/fnins.2013.00267.

Green, D.M., and Swets, J.A. (1974). Signal detection theory and psychophysics.

Haggard, P. (2017). Sense of agency in the human brain. Nat Rev Neurosci 18, 196–207, 10.1038/nrn.2017.14.

Hillyard, S.A., Squires, K.C., Bauer, J.W., and Lindsay, P.H. (1971). Evoked potential correlates of auditory signal detection. Science 172, 1357–1360, 10.1126/science.172.3990.1357.

Horvath, J. (2015). Action-related auditory ERP attenuation: Paradigms and hypotheses. Brain Res 1626, 54–65, 10.1016/j.brainres.2015.03.038.

Hughes, G., and Waszak, F. (2011). ERP correlates of action effect prediction and visual sensory attenuation in voluntary action. Neuroimage 56, 1632–1640, 10.1016/j.neuroimage.2011.02.057.

Jones, S.R., Pritchett, D.L., Stufflebeam, S.M., Hämäläinen, M., and Moore, C.I. (2007). Neural correlates of tactile detection: a combined magnetoencephalography and biophysically based computational modeling study. Journal of Neuroscience 27, 10751–10764, 10.1523/JNEUROSCI.0482-07.2007.

Kilteni, K., and Ehrsson, H.H. (2020). Functional connectivity between the cerebellum and somatosensory areas implements the attenuation of self-generated touch. Journal of Neuroscience 40, 894–906, 10.1523/JNEUROSCI.1732-19.2019.

Lee, C.-C., and Middlebrooks, J.C. (2011). Auditory cortex spatial sensitivity sharpens during task performance. Nature neuroscience 14, 108, 10.1038/nn.2713.

Morillon, B., Schroeder, C.E., and Wyart, V. (2014). Motor contributions to the temporal precision of auditory attention. Nat Commun 5, 5255, 10.1038/ncomms6255.

Morillon, B., Schroeder, C.E., Wyart, V., and Arnal, L.H. (2016). Temporal Prediction in lieu of Periodic Stimulation. J Neurosci 36, 2342–2347, 10.1523/JNEUROSCI.0836-15.2016.

Nelson, A., Schneider, D.M., Takatoh, J., Sakurai, K., Wang, F., and Mooney, R. (2013). A circuit for motor cortical modulation of auditory cortical activity. J Neurosci 33, 14342–14353, 10.1523/JNEUROSCI.2275-13.2013.

Neukirch, M., Hegerl, U., Kötitz, R., Dorn, H., Gallinat, U., and Herrmann, W. (2002). Comparison of the amplitude/intensity function of the auditory evoked N1m and N1 components. Neuropsychobiology 45, 41–48, 10.1159/000048672.

Palmer, C.E., Davare, M., and Kilner, J.M. (2016). Physiological and perceptual sensory attenuation have different underlying neurophysiological correlates. Journal of neuroscience 36, 10803–10812, 10.1523/JNEUROSCI.1694-16.2016.

Reznik, D., Henkin, Y., Levy, O., and Mukamel, R. (2015a). Perceived loudness of self-generated sounds is differentially modified by expected sound intensity. PLoS One 10, e0127651, 10.1371/journal.pone.0127651.

Reznik, D., Henkin, Y., Schadel, N., and Mukamel, R. (2014). Lateralized enhancement of auditory cortex activity and increased sensitivity to self-generated sounds. Nat Commun 5, 4059, 10.1038/ncomms5059.

Reznik, D., and Mukamel, R. (2019). Motor output, neural states and auditory perception. Neurosci Biobehav Rev 96, 116–126, 10.1016/j.neubiorev.2018.10.021.

Reznik, D., Ossmy, O., and Mukamel, R. (2015b). Enhanced auditory evoked activity to self-generated sounds is mediated by primary and supplementary motor cortices. J Neurosci 35, 2173–2180, 10.1523/JNEUROSCI.3723-14.2015.

Saleem, A.B., Ayaz, A., Jeffery, K.J., Harris, K.D., and Carandini, M. (2013). Integration of visual motion and locomotion in mouse visual cortex. Nat Neurosci 16, 1864–1869, 10.1038/nn.3567.

Saupe, K., Widmann, A., Trujillo-Barreto, N.J., and Schroger, E. (2013). Sensorial suppression of self-generated sounds and its dependence on attention. Int J Psychophysiol 90, 300–310, 10.1016/j.ijpsycho.2013.09.006.

Schneider, D.M., Nelson, A., and Mooney, R. (2014). A synaptic and circuit basis for corollary discharge in the auditory cortex. Nature 513, 189–194, 10.1038/nature13724.

Schroeder, C.E., Lindsley, R.W., Specht, C., Marcovici, A., Smiley, J.F., and Javitt, D.C. (2001). Somatosensory input to auditory association cortex in the macaque monkey. Journal of neurophysiology 85, 1322–1327, 10.1523/JNEUROSCI.1694-16.2016.

Shergill, S.S., Samson, G., Bays, P.M., Frith, C.D., and Wolpert, D.M. (2005). Evidence for sensory prediction deficits in schizophrenia. Am J Psychiatry 162, 2384–2386, 10.1176/appi.ajp.162.12.2384.

Sperry, R.W. (1950). Neural basis of the spontaneous optokinetic response produced by visual inversion. J Comp Physiol Psychol 43, 482–489, 10.1037/h0055479.

Tal, I., and Abeles, M. (2013). Cleaning MEG artifacts using external cues. Journal of Neuroscience Methods 217, 31–38, 10.1016/j.jneumeth.2013.04.002.

Timm, J., SanMiguel, I., Saupe, K., and Schröger, E. (2013). The N1-suppression effect for self-initiated sounds is independent of attention. BMC neuroscience 14, 2, 10.1186/1471-2202-14-2.

Uusitalo, M.A., and Ilmoniemi, R.J. (1997). Signal-space projection method for separating MEG or EEG into components. Medical and Biological Engineering and Computing 35, 135–140, 10.1007/BF02534144.

Von Holst, E. (1954). Relations between the central nervous system and the peripheral organs. British Journal of Animal Behaviour, 10.1016/S0950-5601(54)80044-X.

Walsh, L.D., Moseley, G.L., Taylor, J.L., and Gandevia, S.C. (2011). Proprioceptive signals contribute to the sense of body ownership. The Journal of physiology 589, 3009–3021, 10.1113/jphysiol.2011.204941.

Weiss, C., Herwig, A., and Schutz-Bosbach, S. (2011). The self in action effects: selective attenuation of self-generated sounds. Cognition 121, 207–218, 10.1016/j.cognition.2011.06.011.

Zhou, M., Liang, F., Xiong, X.R., Li, L., Li, H., Xiao, Z., Tao, H.W., and Zhang, L.I. (2014). Scaling down of balanced excitation and inhibition by active behavioral states in auditory cortex. Nat Neurosci 17, 841–850, 10.1038/nn.3701.

